# Affinity series of genetically encoded high sensitivity Förster Resonance Energy Transfer sensors for sucrose

**DOI:** 10.1101/2020.11.22.393041

**Authors:** Mayuri Sadoine, Mira Reger, Ka Man Wong, Wolf B. Frommer

**Affiliations:** Institute for Molecular Physiology, Heinrich Heine University Düsseldorf; Department of Plant Biology, Carnegie Institution for Science, Stanford, CA 94305, USA; Institute for transformative Biomolecules, ITbM, Nagoya University, Nagoya, Japan

**Keywords:** sucrose, dynamic range, FLIPsuc nanosensor, specificity, plants

## Abstract

Genetically encoded fluorescent sugar sensors are valuable tools for the discovery of transporters and for quantitative monitoring of sugar steady-state levels in intact tissues. Genetically encoded Förster Resonance Energy Transfer sensors for glucose have been designed and optimized extensively, and a full series of affinity mutants is available for *in vivo* studies. However, to date, only a single improved sensor FLIPsuc-90µΔ1 with a K_m_ for sucrose of ∼90 µM is available for sucrose monitoring. This sucrose sensor was engineered on the basis of an *Agrobacterium tumefaciens* sugar binding protein. Here, we took a two-step approach to first systematically improve the dynamic range of the FLIPsuc nanosensor and then expand the detection range from micromolar to millimolar sucrose concentrations by mutating a key residue in the binding site. The resulting series of sucrose sensors may allow systematic investigation of sucrose transporter candidates and comprehensive *in vivo* analyses of sucrose concentration in plants. Since FLIPsuc-90µ also detects trehalose in animal cells, the new series of sensors can be used to investigate trehalose transporter candidates and monitor trehalose steady-state levels *in vivo* as well.

---

Most methods for the analysis of metabolites rely on destructive methodology and therefore typically lack spatial and temporal resolution. Genetically encoded fluorescent biosensors can overcome these limitations by enabling *in vivo* analyses and by providing rapid (millisecond range) time resolution as well as genetically-based spatial resolution^1^. When no targeting signal is added, these biosensors enable analysis of cytosolic analyte levels. Alternatively, they can be targeted to various compartments such as the endoplasmic reticulum (ER), plasma membrane and nucleus or even attached to a specific protein to provide information specifically from these specific micro-environments^2,3^. The original concept for such sensors was developed for measuring calcium dynamics by making use of conformational rearrangements in calmodulin, or calmodulin and a fused calmodulin binding domain and two spectral variants of the Green Fluorescent Protein that show Förster Resonance Energy Transfer (FRET)^4,5^. The concept was expanded to metabolites by making use of conformational rearrangements in bacterial periplasmic maltose, glucose and ribose binding proteins^6–8^. Since no *bona fide* sucrose binding protein was known to bind sucrose, a candidate protein from *Agrobacterium tumefaciens* was used to engineer a sucrose sensor^9^. However, the initial sucrose sensor had a poor selectivity for sucrose and instead efficiently recognized glucose and maltose. Sequence alignments and structure-guided mutagenesis enabled the generation of a highly selective sensor for sucrose with an affinity of ∼90 µM, termed FLIPsuc-90µ for Fluorescent Indicator Protein for sucrose with an affinity of ∼90 µM. The variant FLIPsuc-90µΔ1, which carried a deletion in one of the linkers showed improved sucrose-induced ratio changes and a larger dynamic range^9^. FLIPsuc-90µΔ1 also recognized trehalose and was successfully used to monitor trehalose transport activity in mammalian cells^10^. Arabidopsis plants, stably transformed with the sucrose sensor, were used to study reversible sucrose accumulation in intact Arabidopsis roots^11,12^. The insensitivity of sucrose accumulation in roots towards protonophores lead to the hypothesis that plant roots express a yet unknown sucrose uniport system^12^. The sucrose sensor successfully used to identify a new class of sugar transporters, the SWEETs, which likely function as sucrose uniporters and which play eminent roles in carbon allocation in Arabidopsis, rice and maize as well as in pathogen susceptibility^13–20^. An efficient analysis of sucrose levels in plant tissues using sucrose sensors was however hampered by both the comparatively low dynamic range and the low signal-to-noise of this initial sucrose sensor FLIPsuc-90µ Δ1. Here, we systematically improved the dynamic range of the FLIPsuc nanosensor by varying linkers between binding protein and fluorescent proteins (FPs) and by exchanging FPs and by mutating the predicted binding site to expand the detection range from low micromolar to millimolar.

## RESULTS AND DISCUSSION

### Strategy to optimize the FLIPsuc nanosensor

The sucrose nanosensor FLIPsuc-Δ1^eCFP-eYFP^ served as basis for the generation of the new sensor series and affinity variants. FLIPsuc-Δ1^eCFP-eYFP^ is composed of the rhizobial sucrose binding protein “ThuE” from *Agrobacterium tumefaciens*, sandwiched between the N-terminal enhanced cyan FP (eCFP) and the C-terminal enhanced yellow FP (eYFP) (Fig. 1)^9^. *At*ThuE had been genetically modified to bind sucrose with high specificity and to reduce the affinity for other sugars such as maltose and glucose. To enhance the sensitivity, the flanking amino acid linkers connecting the FPs to the ThuE binding-domain were shortened in FLIPsuc-90µΔ1^eCFP-eYFP^ compared to the original FLIPsuc-90µ^eCFP-eYFP^ (ref. ^9^). The optimization of the dynamic range and signal-to-noise ratio requires increases of the eYFP/eCFP emission ratio change (ΔR) in response to ligand binding. We therefore sequentially altered key parameters known to be critical in optical sensor optimization such as FP combination, FP dimerization, linker length, and modifications of the binding region by site-directed mutagenesis. Here, we employed stepwise modifications to generate a high sensitivity affinity series of sensors for sucrose (Table 1).

**Table 1:**
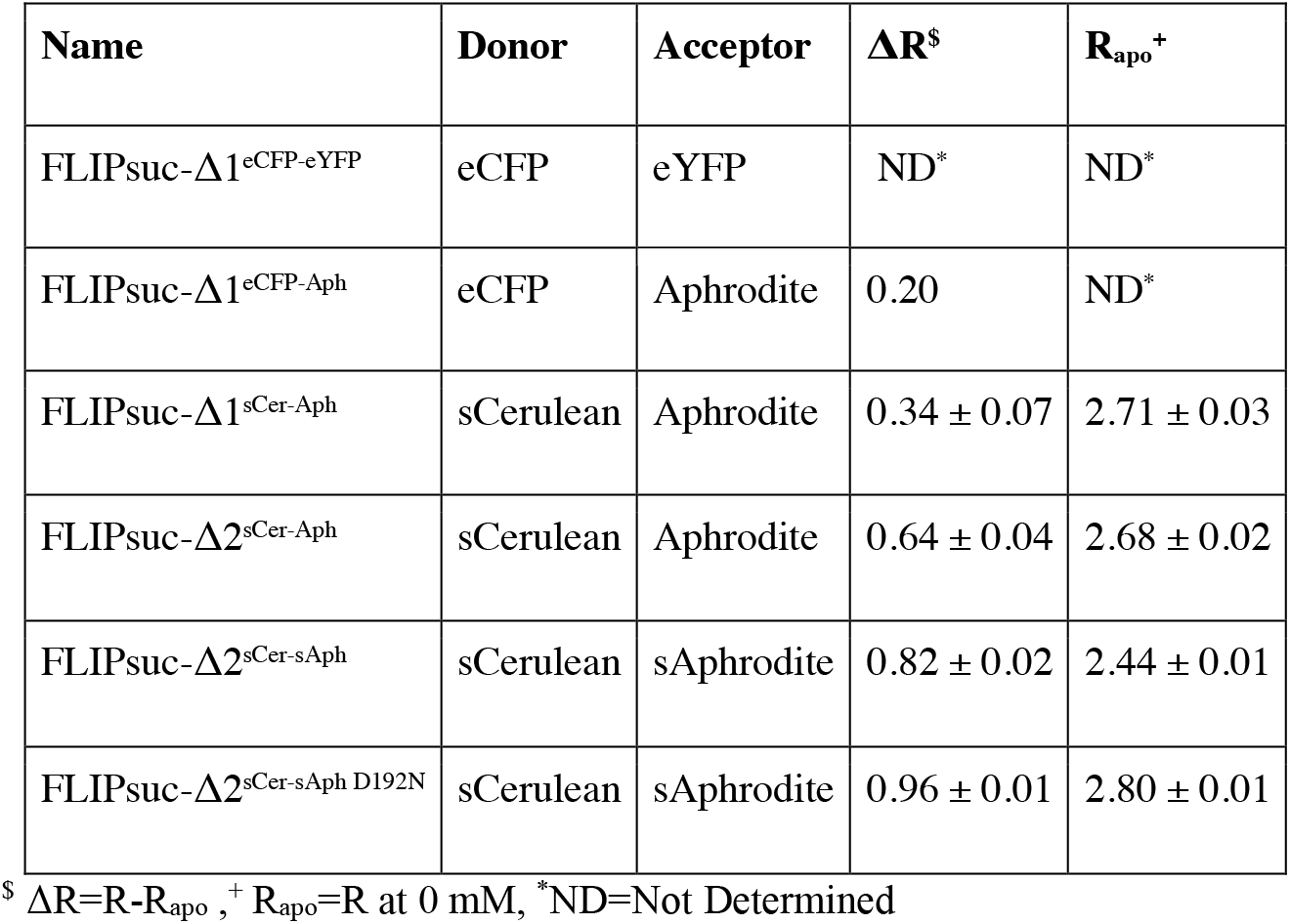
Nomenclature and properties of the step-by-step modified FLIP-Suc.

| Name | Donor | Acceptor | $\Delta R^{\S}$ | $R_{apo}^{+}$ |
| --- | --- | --- | --- | --- |
| FLIPsuc-$\Delta 1^{eCFP-eYFP}$ | eCFP | eYFP | ND* | ND* |
| FLIPsuc-$\Delta 1^{eCFP-Aph}$ | eCFP | Aphrodite | 0.20 | ND* |
| FLIPsuc-$\Delta 1^{sCer-Aph}$ | sCerulean | Aphrodite | $0.34 \pm 0.07$ | $2.71 \pm 0.03$ |
| FLIPsuc-$\Delta 2^{sCer-Aph}$ | sCerulean | Aphrodite | $0.64 \pm 0.04$ | $2.68 \pm 0.02$ |
| FLIPsuc-$\Delta 2^{sCer-sAph}$ | sCerulean | sAphrodite | $0.82 \pm 0.02$ | $2.44 \pm 0.01$ |
| FLIPsuc-$\Delta 2^{sCer-sAph D192N}$ | sCerulean | sAphrodite | $0.96 \pm 0.01$ | $2.80 \pm 0.01$ |

$\S \Delta R = R - R_{apo}$ , $^{+} R_{apo} = R$ at 0 mM, \*ND=Not Determined

**Figure 1:**
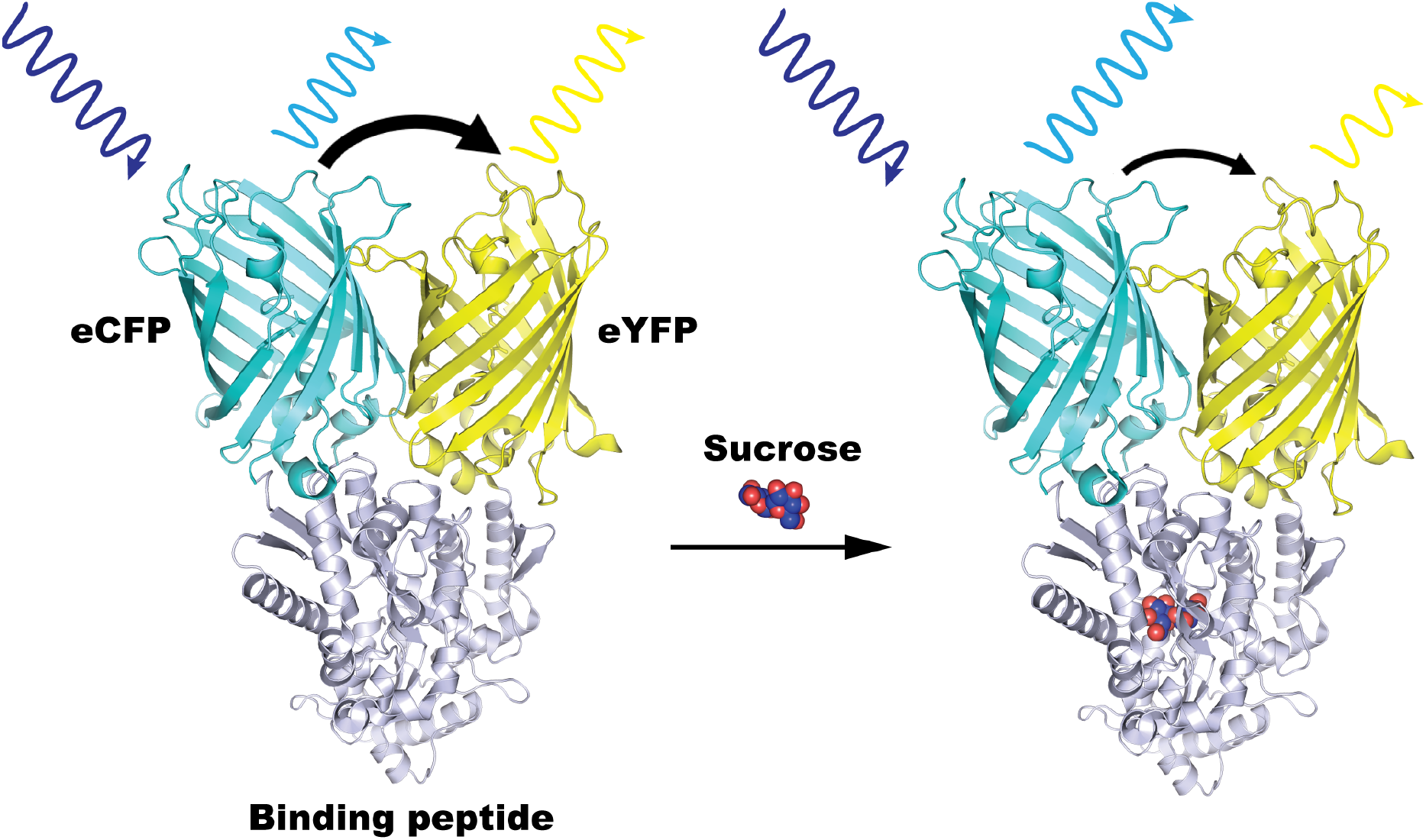
Model of the optical sucrose biosensor FLIPsuc-Δ1^eCFP-eYFP^ (ref. ^9^). The sensor is built by fusing three polypeptides: A FRET donor FP, eCFP, (cyan), a sandwiched binding peptide, ThuE, (grey) and a FRET acceptor FP, eYFP (yellow). The sensor mechanism is displayed with an unbound state associated to high eYFP/eCFP emission ratio (left) and sucrose bound state associated with low eYFP/eCFP emission ratio (right). The binding of a sucrose molecule drives conformational rearrangements leading to a decrease in the eYFP/eCFP emission ratio in the equivalent response

### Testing effect of altered FP combinations

It had been shown previously that exchanging the FPs in nanosensors impacts sensor properties^21^. Therefore, we first replaced the N-terminal acceptor FP with Aphrodite (Aph), a codon-modified version of Venus. The resulting sensor FLIP-suc-Δ1^eCFP-Aph^, which also contains a shortened linker, showed only a modest ratio change of 0.2 (Table 1 and Table S1). We next replaced the eCFP in FLIPsuc-Δ1^eCFP-Aph^ with a variety of cyan emitting FPs (Table S1), while maintaining Aph as C-terminal FP, but none of the sensor variants showed an improved ratio change compared to FLIPsuc-Δ1^eCFP-Aph^. However, when a modified, so called “sticky” version of Cerulean (sCer)^22^, was introduced as donor FP, the ratio change increased by ∼50 % (Table 1 and Fig. 2A) in FLIPsuc-Δ1^sCer-Aph^. The sCer variant contains two point mutations, a substitution of serine at position 208 in the dimerization interface with phenylalanine (S208F), and a substitution of valine at position 224 close to the chromophore with leucine (V224L), respectively (Fig. 2A-C). It was found that the substitutions S208F and V224L yielded the biggest improvement of the ratio change of all fluorophore pairs tested, possibly due to the rearrangement of the FP-dimerization interface and stabilization of the chromophore orientation inside the FP barrel^23,24^. Substitutions S208F and V224L were shown to substantially improve the response to sucrose of FLIPsuc-Δ1^sCer-Aph^, while maintaining high selectivity for sucrose (Fig. 2E and 2D).

**Figure 2:**
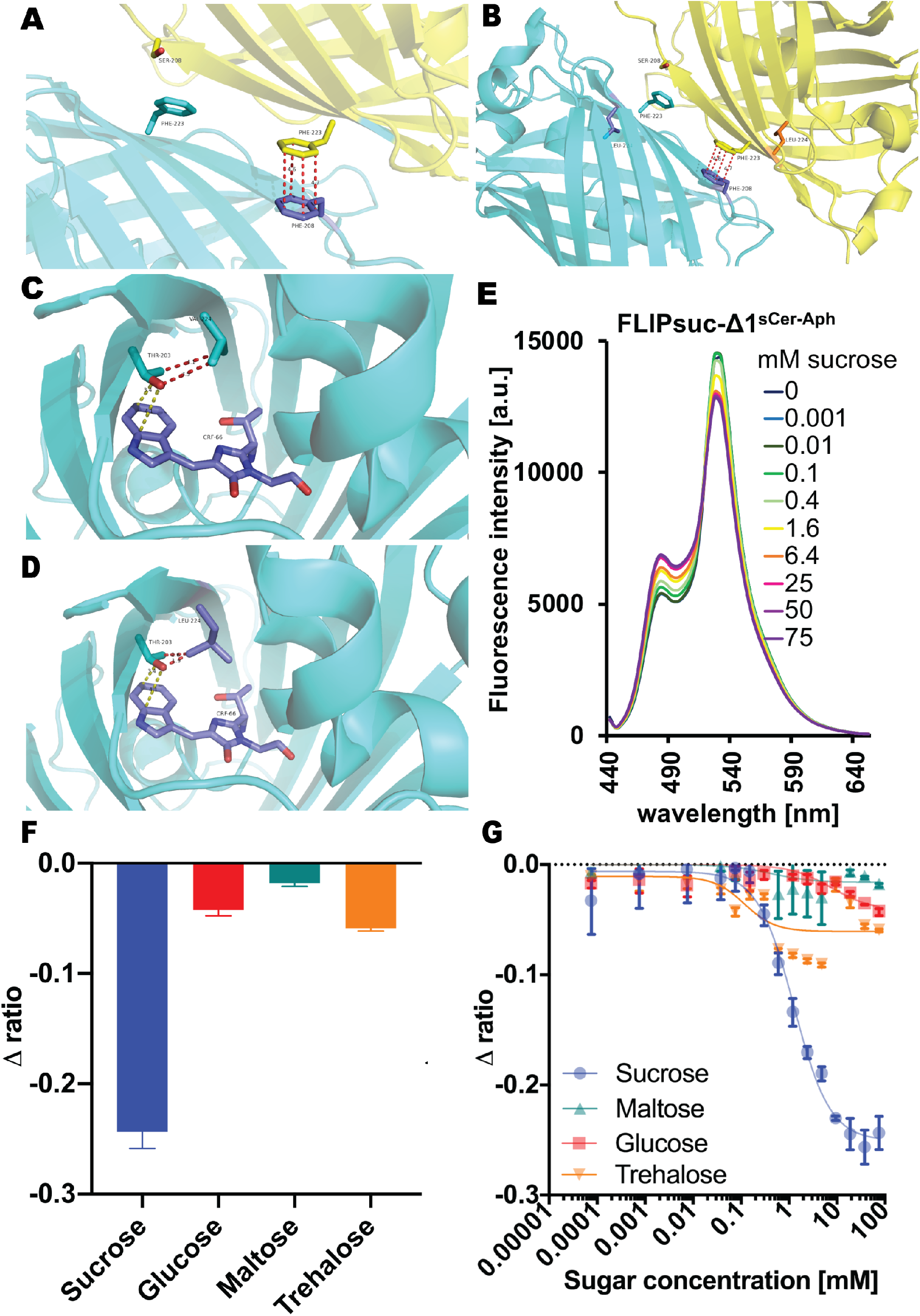
Combination of sCerulean and Aphrodite in FLIPsuc-Δ1^sCer-Aph^ led to an increase in ratio change. Possible rearrangement of the dimerization interface due to substitutions S208F and V224L in Cerulean donor FP (A-D) in FLIPsuc-Δ1sCer-Aph affect R_apo_ and ratio change upon sucrose binding (E) while maintaining high sucrose specificity (F, G).

### Optimization of the length of FP linkers

To further enhance the dynamic range, FLIPsuc-90µΔ1^sCer-Aph^ was subjected to extensive linker modifications (Fig. S1). FPs generally contain terminal linker regions that are not essential for folding and fluorescence of the FP and can thus be modified to decrease flexibility and maximize dipole coupling^25^. It had been shown previously, that the linker length impacts the emission acceptor/donor ratio of the apo state (R_apo_) and the dynamic range^9,26^. Therefore, we systematically truncated the linker residues connecting sCer with ThuE (termed N-terminal linker) and the linker residues connecting ThuE with Aph (termed C-terminal linker). The N-terminal linker composition was systematically decreased from 13 to 2 amino acids and the C-terminal linker decreased from 11 to 2 amino acids (Fig. S1A). The linker combination of 12 N-terminal residues and 5 C-terminal residues (L10/R4) yielded the best performing sensor, with a ratio change of about 0.7 (Fig. S1B) in FLIPsuc-Δ2^sCer-Aph^.

### Modification of the acceptor FP and FP dimerization

Modification of the C-terminal acceptor FP in FLIPsuc-Δ2^sCer-Aph^ may affect the ratio change^21^. Thus, we further evaluated the effects of FP dimerization capability of the C-terminal Aph with sCer in FLIPsuc-Δ2^sCer-Aph^. To increase the effects of FP-dimerization and thus improve the R_apo_ and ratio change upon sucrose binding ΔR, the same substitutions compared to sCer, namely S208F and V224L, were introduced into Aph (Fig. 3 and Table 2). A sensor variant with the single substitution Aph^S208F^ displayed a substantial increase in R_apo_ of ∼90% compared to FLIPsuc-Δ2^sCer-Aph^ with the non-mutated Aph, but the ratio change upon sucrose binding was strongly reduced (Fig. 3A and Table 2).. Similar results were obtained with the double substitutions S208F and V224L (Table 2). Interestingly, the sensor with Aph^V224L^ showed an increase to 0.8 in ΔR while maintaining high selectivity towards sucrose (Fig. 3B-D and Table 2). We postulate that the substitution V224L stabilizes the Aph-chromophore (Fig. 3E and 3F). Thus, the combination of sCer with the single substitution V224L in of Aph in the newly designed FLIPsuc-Δ2^sCer-sAph^ yielded the best ratio change upon ligand treatment.

**Table 2.** Optimization of FP dimerization properties by site-directed mutagenesis in acceptor FP.

| Name | FP<br>N-terminus | FP C-terminus | $\Delta R^{\S}$ | $R_{apo}^{+}$ |
| --- | --- | --- | --- | --- |
| FLIPsuc-$\Delta 2^{sCer-Aph}$ | sCer | Aph | $0.64 \pm 0.04$ | $2.68 \pm 0.02$ |
| FLIPsuc-$\Delta 2^{sCer-Aph S208F}$ | sCer | Aph S208F | $0.34 \pm 0.04$ | $5.27 \pm 0.04$ |
| FLIPsuc-$\Delta 2^{sCer-Aph S208F/V224L}$ | sCer | Aph S208F/V224L | $0.65 \pm 0.04$ | $6.81 \pm 0.06$ |
| FLIPsuc-$\Delta 2^{sCer-sAph}$ | sCer | Aph V224L | $0.82 \pm 0.02$ | $2.44 \pm 0.01$ |

$\S$ Normalized ratio= $R - R_{apo}$ , $^{+} R_{apo} = R$ at 0mM

**Figure 3:**
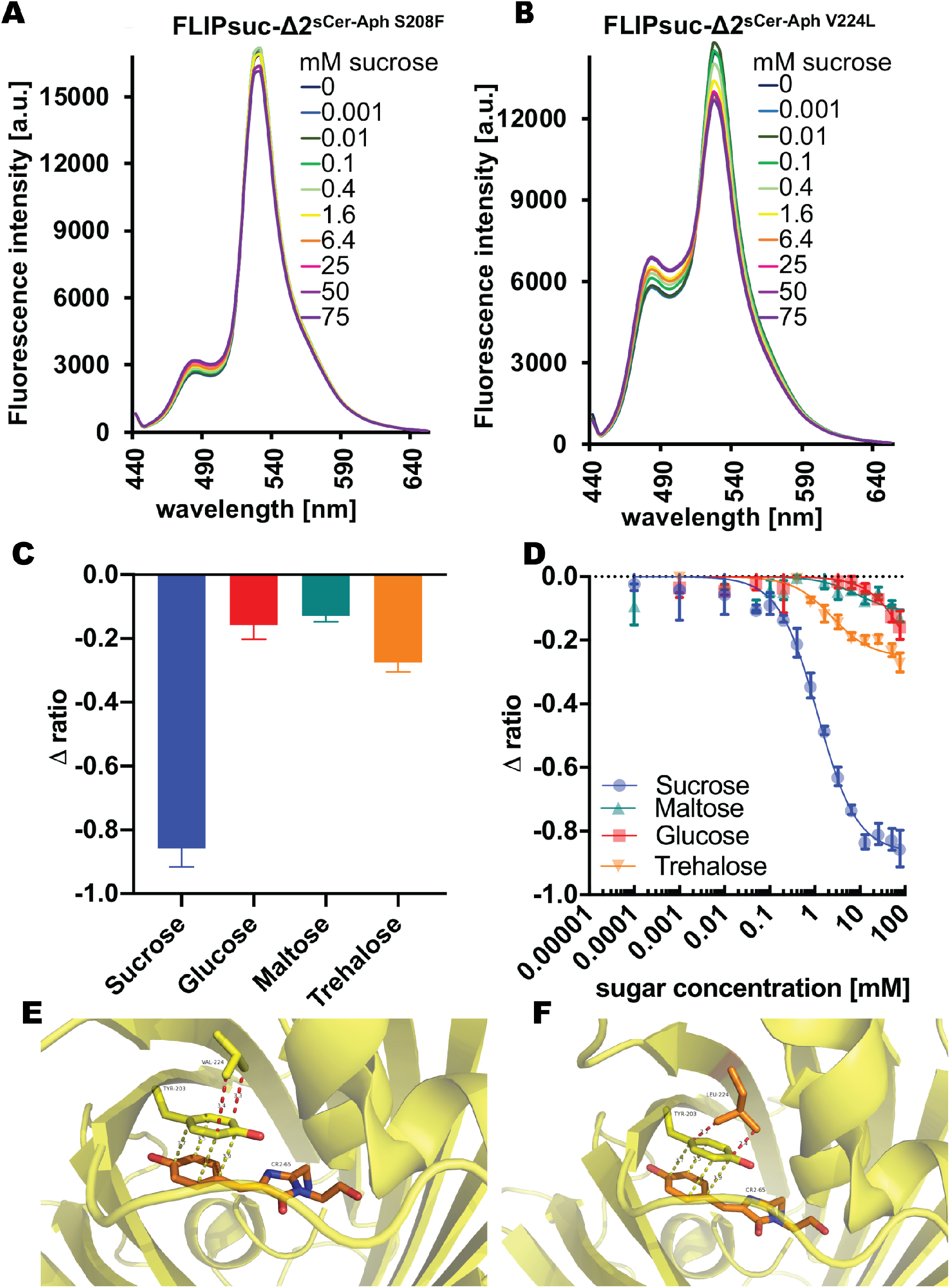
Substitution V224L in Aphrodite of FLIPsuc-Δ2^sCer-sAph^ improves FP dimerization and increases the emission ratio change. In Aphrodite, single substitution S208F decreases ratio change (A) while V224L increases ratio change (B). The combination of double substitutions S208F and V224L in Cerulean and single substitution V224L in Aphrodite led to substantial improvement of the sensor while maintaining high sucrose specificity (C, D). This is most likely because V224L in Aphrodite (E, F) leads to rearrangement of the dimerization interface.

### Substitution D192N in ThuE binding region

The original sensor FLIPsuc-90μ carried a N192D substitution relative to the published sequence from *At*ThuE, which is located outside the binding pocket^9^. Whether this substitution is due to natural variation between strains or occurred during the initial cloning remains unknown. To test whether the substitution is relevant for sensor properties, site-directed mutagenesis on FLIPsuc-Δ2^sCer-sAph^ was performed to revert the sequence of the published ThuE sequence. After introducing the substitution D192N in ThuE of FLIPsuc-Δ2^sCer-sAph D192N^, we found that the sensor showed an increase in ΔR to about 0.95 (Table 1 and Fig. 4A and 4B). Since the substitution was introduced in a small beta sheet structure of the binding domain in vicinity of the binding interface while yet outside the actual binding pocket, we investigated whether the selectivity or the affinity towards sucrose were affected (Fig. 4C and 4D). Indeed, D192N introduction in FLIPsuc-90µΔ2^sCer-sAph^ led to an increase in the affinity to 65 µM for sucrose (Table 3). Surprisingly, FLIPsuc-Δ2^sCer-sAph D192N^ was also more selective towards sucrose (Fig. 4E and 4F). We have designed and characterized the affinity series of FLIPsuc with this nanosensor, FLIPsuc-Δ2^sCer-sAph D192N^.

**Table 3.** FLIPsuc-Δ2^sCer-sAph D192N^ affinity mutant series.

| Name | Mutation<br>in ThuE | Affinity<br>[$\mu M$] | Detection<br>range [$\mu M$] |
| --- | --- | --- | --- |
| FLIPsuc-$65\mu \Delta 2^{sCer-sAph D192N}$ | W283A | $65 \pm 3$ | 7-585 |
| FLIPsuc-$90\mu \Delta 2^{sCer-sAph D192N}$ | W283G | $87 \pm 3$ | 10-783 |
| FLIPsuc-$450\mu \Delta 2^{sCer-sAph D192N}$ | W283C | $454 \pm 20$ | 50-4086 |
| FLIPsuc-$1.1m \Delta 2^{sCer-sAph D192N}$ | W283I | $1091 \pm 48$ | 121-9819 |
| FLIPsuc-$2.4m \Delta 2^{sCer-sAph D192N}$ | W283V | $2414 \pm 136$ | 268-21726 |

**Figure 4:**
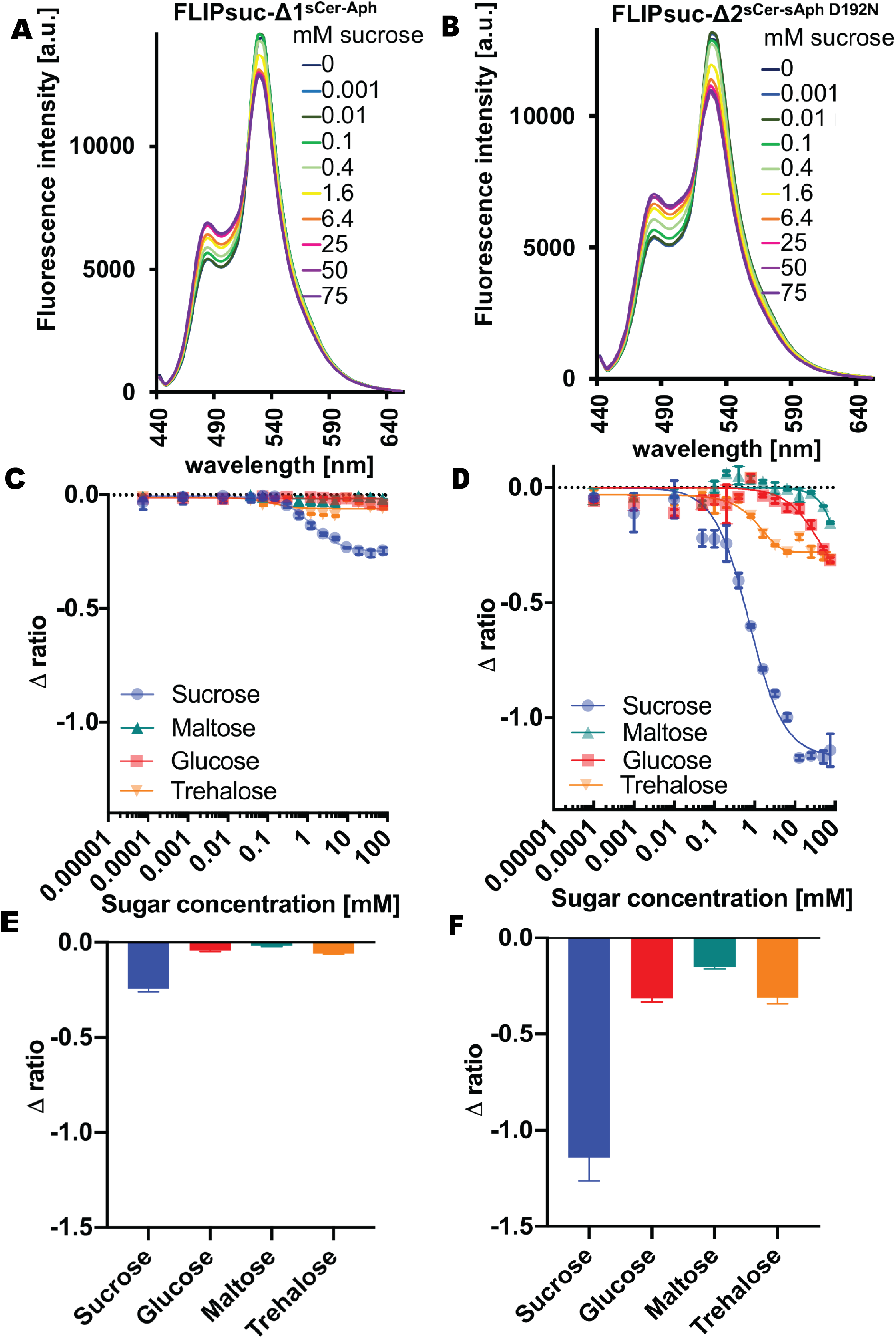
Substitution D192N in FLIPsuc-Δ2^sCer-sAph^ improves the sensor response. Single substitution D192N in ThuE binding protein increases ratio change from FLIPsuc-Δ1^sCer-Aph^ (A-F). The substitution in ThuE binding region led to a slight change in the sensor affinity while maintain high sucrose specificity (C and F).

### Generation of a high sensitivity affinity series of FLIPsuc nanosensors

In the absence of reference information on the actual sucrose levels in the different compartments and cells in a plant, and given the limited detection range of a sensor of about two orders of magnitude (∼6.5 µM to 650 µM for FLIPsuc-65µΔ2^sCer-sAph D192N^), *in planta* analyses will require a series of sucrose sensors with different affinities. We had shown previously that the binding pocket of ThuE can be altered to improve selectivity for sucrose^9^. Exchange of W283A had been shown to have the highest effect on ligand binding specificity (Fig. 5A) and led to the design of FLIPsuc-90µ (ref.^9^). Here, we systematically substituted W283 in the nanosensor to generate a set of affinity mutants ranging from micromolar to millimolar. We substituted the original W283 with arginine, glycine, cysteine, leucine, isoleucine and valine, in both FLIPsuc-Δ2^sCer-Aph^ and FLIPsuc-Δ2^sCer-sAph D192N^ (Tables 3 and S4, Fig. 5B-5D). We found that FLIPsuc-Δ2^sCer-sAph D192N^ derivates covered a broad coverage of concentrations with K_d_ of 65 μM, 90 μM, 450 μM, 1.1 mM and 2.4 mM for the substitutions W283A, W283G, W283C, W283L, W283I and W283V, respectively (Fig. 5D). Since the detection range is defined as the region between 10 % and 90 % of the response of a sensor, FLIPsuc-Δ2^sCer-sAph D192N^ affinity series constitutes thus a set of sucrose nanosensors with a broad detection range between 7 µM and 21.7 mM.

**Figure 5:**
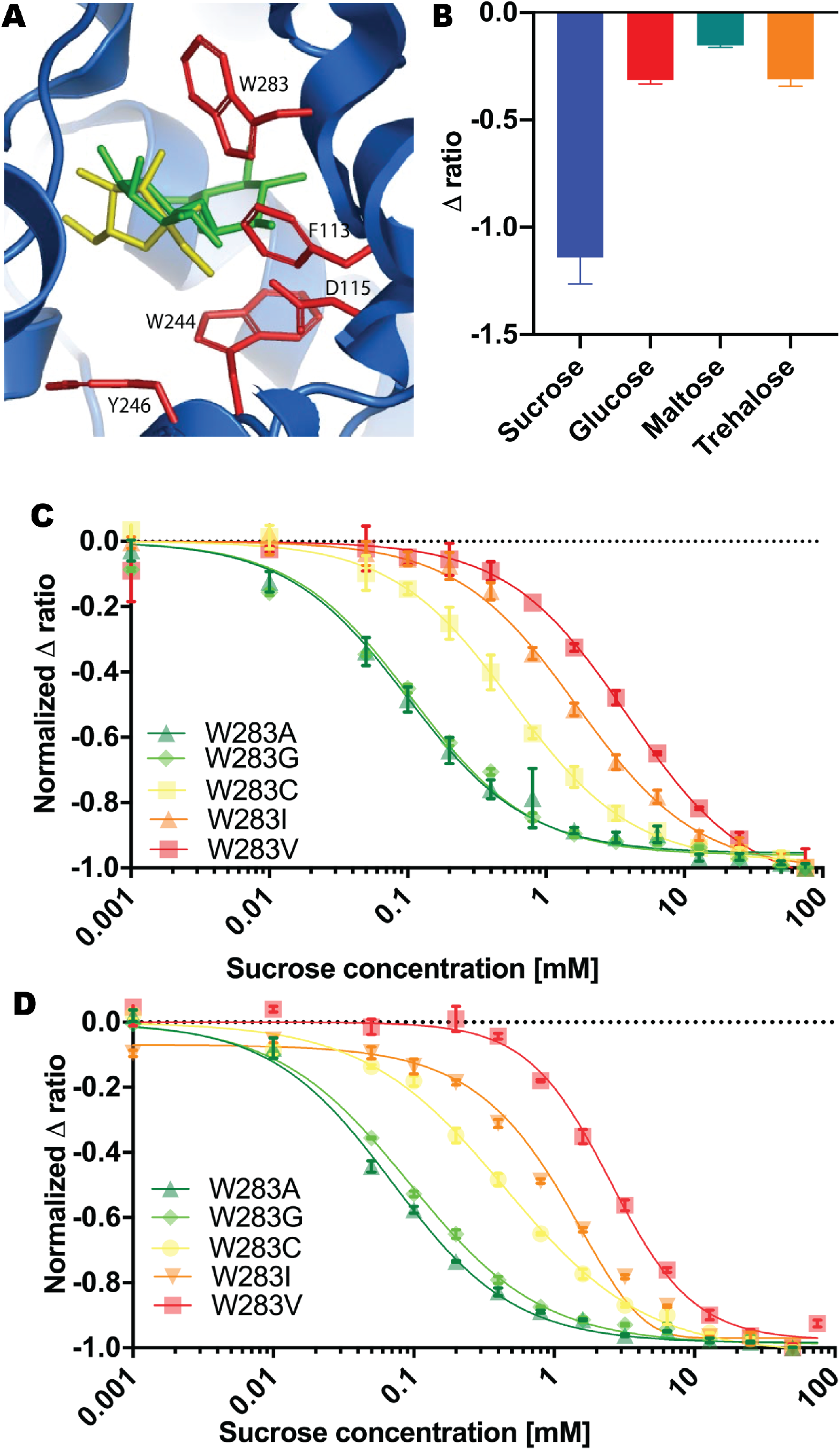
Systemic substitutions of W283 in binding pocket of FLIPsuc change sensor affinity. Systematic substitutions of W283, which directly interacts with the sucrose molecule (A), likely affects the affinity as it was previously found when W283A led to high specificity to sucrose and design of FLIPsuc-90μ (A, B)^9^. Systematic substitutions W283A, W283G, W283C, W283L, W283I and W283V in ThuE of FLIPsuc-Δ2^sCer-Aph^ (C) and FLIPsuc-Δ2^sCer-sAph D192N^ (D) allows the design of a series of affinity mutants with K_m_ from micromolar to millimolar and broad detection range.

## CONCLUSION

Here we present a series of improved sucrose sensors named FLIPsuc-Δ2^sCer-sAph D192N^ with improved dynamic range, signal to noise and a broad detection range. The resulting sucrose sensor series is constituted of highly sucrose-selective mutants with affinities ranging from micromolar to millimolar concentrations for sucrose. This new set of FLIPsuc sensors represents a valuable set of tools for functional characterization of sucrose transporters and for monitoring of wide concentration range of sucrose *in planta* as well as trehalose in animals.

## MATERIAL AND METHODS

### Constructs of FLIPsuc nanosensors

Constructs of pRSET (Invitrogen) carrying S208F, V224L in Cer, S208F, V224L in Aph and D192N, W283A W283G, W283C, W283L, W283I and W283V, respectively in ThuE were designed by site-directed mutations in optical sucrose sensors by using the QuickChange II Site-Directed Mutagenesis (SDM) Kit (Agilent Technologies) following the manufacturer’s instructions. Nanosensor sequences were confirmed by DNA sequencing.

### Nanosensor expression and purifications

Proteins were expressed in BL21(DE3)Gold (Stratagene) and purified as described previously^6^. In brief, cells were grown for 2 days in the dark at 21°C and then harvested by centrifugation, resuspended in 20 mM Tris·Cl, pH 7.9, and disrupted by ultrasonication. His tagged FLIPsuc sensors were purified by affinity chromatography using Ni-NTA bead material (Novagen). Binding to the resin was performed in batch at 4 °C for 4 h, and followed by washing steps with 20 mM Tris·HCl and 20 mM Tris·HCl containing 20 mM imidazole at pH 7.9. Elution was performed with 200 mM imidazole in Tris·HCl, pH 7.9.

### In vitro characterization of FLIPsuc and binding assays

*In vitro* characterization of the sucrose sensors was performed in transparent 96-well microtiter plates with flat bottom wells on a Safire (Tecan) fluorimeter. Before each measurement, the sensor solutions were diluted in 20 mM MOPS buffer pH 8.0. Similar amount of sensors were ensured by measuring the Aphrodite fluorescence emission at λ_em_ 530 nm via direct excitation at λ_ex_ of 505 nm. Binding assays were performed by using sucrose or glucose, maltose, trehalose in 20 mM MOPS pH 8.0 added to final concentrations from 0 mM to 75 mM in wells. Sensor-ligand titration curves and ligand specificity experiments were conducted as follow. Cer and sCer of the sensors were excited at λ_ex_ 420 nm and fluorescent emissions were recorded over the range from λ_em_ 440 nm to λ_em_ 650 nm in Δλ 5 nm steps for full emission spectrum analyses. Ligand titration curves and ligand specificity analyses were performed on a Safire (Tecan) fluorimeter 2 days after protein purifications which ensure that all FPs are completely matured at the time of measurement. Emission ratio point measurements were performed with Cer excited at 428 nm, and recording emission maxima of Cer/sCer and Aph/sAph at 485 and 530 nm, respectively with a bandwidth of 12 nm. The change in the relative emission ratio of the two fluorophores was determined by Aph/Cer emission intensity ratio. Data from emission ratio measurements were processed using Microsoft Excel 2010. The fluorescence emission intensity ratios before adding ligand (R_apo_) and with increasing ligand concentration (R_n_) were calculated by dividing the fluorescence intensity of FRET (λ_ex_ 420 nm, λ_em_ 530 nm) by the intensity of Cer (λ_ex_ 420 nm, λ_em_ 480 nm/ 484 nm/ 485 nm). Calculation of the ratios R and ΔR_n_ however was performed with the maximum of Cer emission. Ratio changes upon ligand binding (ΔR_n_) were calculated as the actual FRET response of the sensor:

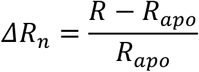

Analysis and visualization of the FRET data was performed using the scientific software GraphPad Prism 6.01. Data were entered in mean and standard deviation of R or ΔR_n_ of three technical replicates. Using the change in ratio upon ligand binding, the K_d_ of each sensor was determined by fitting ligand titration curves to a non linear one-site specific binding with Hill slope and a constant: Y = B_max_ (X^h^)/(K_m_^h^+X^h^) + a, with B_max_ being the maximum specific binding, K_m_ the Michaelis constant, h the Hill slope which describes the cooperativity of the ligand binding and a constant added to the equation. Analyses were performed with at least three independent protein preparations.

### Homology Modeling of proteins

Homology models were created only of the fluorophores Cer (PDB ID: 2Q57), Venus (PDB ID: 1MYW) and the dimerized GFP (PDB ID: 1GFL). Modeling was performed using the molecular visualization system PyMOL 1.7.2 (Schrödinger). The crystal structures of the green fluorescent protein GFP (1GFL), yellow fluorescent protein Venus (1MYW) and cyan fluorescent protein Cerulean (2Q57) were obtained from the RCSB Protein Data Base (PDB). 3D analysis of protein structures and simulation of point mutants was performed using the molecular visualization system PyMOL 1.7.2 (Schrödinger). AtThuE was modelized by using Protean3D.

## ASSOCIATED CONTENT

## AUTHOR INFORMATION

### Author Contributions

BH, MR, KMW have performed experiments, BH, MR, KMW, MS have performed analyzes, MS and WBF have written the manuscript, MS, KMW and WBF have revised the manuscript.

## ACKNOWLEDGMENT

We thank Bi Huei Hou for excellent technical assistance. We also thank Dr. Cindy Ast for fruitful discussion and comments about the manuscript and for generating the FLIPsuc nanosensor three dimensional models.

## ABBREVIATIONS

FRET: Fluorescence Energy Transfer
SWEET: Sugar Will Eventually be Exported Transporter
FP: Fluorescent Protein
ER: Endoplasmic Reticulum
eCFP: enhanced Cyan Fluorescent Protein
eYFP: enhanced Yellow Fluorescent Protein
Aph: Aphrodite
sCer: sticky Cerulean
SDM: Site Directed Mutagenesis
PDB: Protein DataBase
A: Arginine
G: Glycine
C: Cysteine
L: Leucine
I: Isoleucine
V: Valine

## REFERENCES

(1) Okumoto, S., Jones, A., Frommer, W. B. Quantitative Imaging with Fluorescent Biosensors. Annual review of plant biology 2012, 63, 663–706.

(2) Okumoto, S., Looger, L. L., Micheva, K. D., Reimer, R. J., Smith, S. J., Frommer, W. B. Detection of Glutamate Release from Neurons by Genetically Encoded Surface-Displayed FRET Nanosensors. Proceedings of the National Academy of Sciences 2005, 102, 8740–8745.

(3) Takanaga, H., Frommer, W. B. Facilitative Plasma Membrane Transporters Function during ER Transit. The FASEB Journal 2010, 24, 2849–2858.

(4) Romoser, V. A., Hinkle, P. M., Persechini, A. Detection in Living Cells of Ca^2+^-Dependent Changes in the Fluorescence Emission of an Indicator Composed of Two Green Fluorescent Protein Variants Linked by a Calmodulin-Binding Sequence A New Class of Fluorescent Indicators. Journal of biological chemistry 1997, 272, 13270–13274.

(5) Miyawaki, A., Llopis, J., Heim, R., McCaffery, J. M., Adams, J. A., Ikura, M., Tsien, R. Y. Fluorescent Indicators for Ca^+^, Based on Green Fluorescent Proteins and Calmodulin. Nature 1997, 388, 882–887.

(6) Fehr, M., Frommer, W. B., Lalonde, S. Visualization of Maltose Uptake in Living Yeast Cells by Fluorescent Nanosensors. Proceedings of the National Academy of Sciences 2002, 99, 9846–9851.

(7) Lager, I., Fehr, M., Frommer, W. B., Lalonde, S. Development of a Fluorescent Nanosensor for Ribose. FEBS letters 2003, 553, 85–89.

(8) Fehr, M., Lalonde, S., Lager, I., Wolff, M. W., Frommer, W. B. In Vivo Imaging of the Dynamics of Glucose Uptake in the Cytosol of COS-7 Cells by Fluorescent Nanosensors. Journal of Biological Chemistry 2003, 278, 19127–19133.

(9) Lager, I., Looger, L. L., Hilpert, M., Lalonde, S., Frommer, W. B. Conversion of a Putative Agrobacterium Sugar-Binding Protein into a FRET Sensor with High Selectivity for Sucrose. Journal of Biological Chemistry 2006, 281, 30875– 30883.

(10) Kikuta, S., Hou, B.-H., Sato, R., Frommer, W. B., Kikawada, T. FRET Sensor-Based Quantification of Intracellular Trehalose in Mammalian Cells. Bioscience, biotechnology, and biochemistry 2016, 80, 162–165.

(11) Grossmann, G., Guo, W.-J., Ehrhardt, D. W., Frommer, W. B., Sit, R. V., Quake, S. R., Meier, M. The RootChip: An Integrated Microfluidic Chip for Plant Science. The plant cell 2011, 23, 4234–4240.

(12) Chaudhuri, B., Hörmann, F., Lalonde, S., Brady, S. M., Orlando, D. A., Benfey, P., Frommer, W. B. Protonophore-and PH-insensitive Glucose and Sucrose Accumulation Detected by FRET Nanosensors in Arabidopsis Root Tips. The Plant Journal 2008, 56, 948–962.

(13) Chen, L.-Q., Hou, B.-H., Lalonde, S., Takanaga, H., Hartung, M. L., Qu, X.-Q., Guo, W.-J., Kim, J.-G., Underwood, W., Chaudhuri, B., Chermak, D., Antony, G., White, F. F., Somerville, S. C., Mudgett, M. B., Frommer, W. B. Sugar Transporters for Intercellular Exchange and Nutrition of Pathogens. Nature 2010, 468, 527–532.

(14) Chen, L.-Q., Qu, X.-Q., Hou, B.-H., Sosso, D., Osorio, S., Fernie, A. R., Frommer, W. B. Sucrose Efflux Mediated by SWEET Proteins as a Key Step for Phloem Transport. Science 2012, 335, 207–211.

(15) Eom, J.-S., Luo, D., Atienza-Grande, G., Yang, J., Ji, C., Huguet-Tapia, J. C., Char, S. N., Liu, B., Nguyen, H., Schmidt, S. M. Diagnostic Kit for Rice Blight Resistance. Nature biotechnology 2019, 1–8.

(16) Oliva, R., Ji, C., Atienza-Grande, G., Huguet-Tapia, J. C., Perez-Quintero, A., Li, T., Eom, J.-S., Li, C., Nguyen, H., Liu, B. Broad-Spectrum Resistance to Bacterial Blight in Rice Using Genome Editing. Nature biotechnology 2019, 1–7.

(17) Bezrutczyk, M., Hartwig, T., Horshman, M., Char, S. N., Yang, J., Yang, B., Frommer, W. B., Sosso, D. Impaired Phloem Loading in Genome-Edited Triple Knock-out Mutants of SWEET13 Sucrose Transporters. bioRxiv 2017, 197921.

(18) Lin, I. W., Sosso, D., Chen, L.-Q., Gase, K., Kim, S.-G., Kessler, D., Klinkenberg, P. M., Gorder, M. K., Hou, B.-H., Qu, X.-Q. Nectar Secretion Requires Sucrose Phosphate Synthases and the Sugar Transporter SWEET9. Nature 2014, 508, 546–549.

(19) Chen, L.-Q., Lin, I. W., Qu, X.-Q., Sosso, D., McFarlane, H. E., Londoño, A., Samuels, A. L., Frommer, W. B. A Cascade of Sequentially Expressed Sucrose Transporters in the Seed Coat and Endosperm Provides Nutrition for the Arabidopsis Embryo. The Plant Cell 2015, 27, 607–619.

(20) Sosso, D., Luo, D., Li, Q.-B., Sasse, J., Yang, J., Gendrot, G., Suzuki, M., Koch, K. E., McCarty, D. R., Chourey, P. S. Seed Filling in Domesticated Maize and Rice Depends on SWEET-Mediated Hexose Transport. Nature genetics 2015, 47, 1489.

(21) Deuschle, K., Chaudhuri, B., Okumoto, S., Lager, I., Lalonde, S., Frommer, W. B. Rapid Metabolism of Glucose Detected with FRET Glucose Nanosensors in Epidermal Cells and Intact Roots of Arabidopsis RNA-Silencing Mutants. The Plant Cell 2006, 18, 2314–2325.

(22) Lindenburg, L. H., Malisauskas, M., Sips, T., van Oppen, L., Wijnands, S. P., van de Graaf, S. F., Merkx, M. Quantifying Stickiness: Thermodynamic Characterization of Intramolecular Domain Interactions to Guide the Design of Forster Resonance Energy Transfer Sensors. Biochemistry 2014, 53, 6370–6381.

(23) Vinkenborg, J. L., Evers, T. H., Reulen, S. W., Meijer, E. W., Merkx, M. Enhanced Sensitivity of FRET-based Protease Sensors by Redesign of the GFP Dimerization Interface. ChemBioChem 2007, 8, 1119–1121.

(24) Nguyen, A. W., Daugherty, P. S. Evolutionary Optimization of Fluorescent Proteins for Intracellular FRET. Nature biotechnology 2005, 23, 355–360.

(25) Li, X., Zhang, G., Ngo, N., Zhao, X., Kain, S. R., Huang, C.-C. Deletions of the Aequorea Victoria Green Fluorescent Protein Define the Minimal Domain Required for Fluorescence. Journal of Biological Chemistry 1997, 272, 28545–28549.

(26) Deuschle, K., Okumoto, S., Fehr, M., Looger, L. L., Kozhukh, L., Frommer, W. B. Construction and Optimization of a Family of Genetically Encoded Metabolite Sensors by Semirational Protein Engineering. Protein Science 2005, 14, 2304–2314.

