## Supplementary Information for "Affinity series of genetically encoded high sensitivity Förster Resonance Energy Transfer sensors for sucrose"

**Table S1:** Optimization of the donor FP of in FLIPsuc.

| **Name** | **FP N-terminus** | **FP C-terminus** | **ΔR$** |
|---|---|---|---|
| FLIPsuc-90$\mu\Delta1^{eCFP\text{-}Aph}$ | eCFP | Aphrodite | 0.20 |
| FLIPsuc-90$\mu\Delta1^{TFTt9\text{-}Aph}$ | TFPt9 | Aphrodite | 0.03 |
| FLIPsuc-90$\mu\Delta1^{TFP\text{-}Aph}$ | TFP | Aphrodite | 0.15 |
| FLIPsuc-90$\mu\Delta1^{mTrq2\text{-}Aph}$ | mTrq2 | Aphrodite | 0.11 |
| FLIPsuc-90$\mu\Delta1^{sTrq2\text{-}Aph}$ | sTrq2 | Aphrodite | 0.11 |
| FLIPsuc-90$\mu\Delta1^{t7mTrq2t9\text{-}Aph}$ | t7mTrq2t9 | Aphrodite | 0.11 |
| FLIPsuc-90$\mu\Delta1^{Cer\text{-}Aph}$ | Cer | Aphrodite | 0.15 |
| FLIPsuc-90$\mu\Delta1^{mCer\text{-}Aph}$ | mCer | Aphrodite | 0.09 |
| FLIPsuc-90$\mu\Delta1^{sCer\text{-}Aph}$ | sCer | Aphrodite | 0.30 |

$Normalized ratio=$R\text{-}R_{apo}$

**Table S2:** Primers used to revert ThuE sequence and modified Aph FP.

| **Label** | **Sequence** |
|---|---|
| ThuE_D192N_F | GTGGCCATATCGTTGAAACCAACGGCGATATCTCCATCAATAACG |
| ThuE_D192N_R | CGTTATTGATGGAGATATCGCCGTTGGTTTCAACGATATGGCCAC |
| Aph_S208F_F | CTCTCGTACCAAAGCGCGCTCTCCAAGGACCCGAATGAGAAACGG |
| Aph_S208F_R | CCGTTTCTCATTCGGGTCCTTGGAGAGCGCGCTTTGGTACGAGAG |
| Aph_V224L_F | CCACATGGTTCTCCTGGAGTTCCTCACGGCGCGCGGCATATAG |
| Aph_V224L_R | CTATATGCCGCGCGCCGTGAGGAACTCCAGGAGAACCATGTGG |

**Table S3:** Plasmids used in this study.

| Plasmid | Description | Reference |
|---|---|---|
| pRSET_sCer-ThuE-Aph | FLIPsuc-$\Delta 2^{sCer\text{-}Aph}$ | this study |
| pRSET_sCer-ThuE-Aph$^{S208F}$ | FLIPsuc-$\Delta 2^{sCer\text{-}Aph}$ with substitution in S208F in Aphrodite | this study |
| pRSET_sCer-ThuE-Aph$^{V224L}$ | FLIPsuc-$\Delta 2^{sCer\text{-}Aph}$ with substitution V224L in Aphrodite | this study |
| pRSET_sCer-ThuE-Aph$^{S208F/V224L}$ | FLIPsuc-$\Delta 2^{sCer\text{-}Aph}$ with substitution V224L and S208F in Aphrodite | this study |
| pRSET_sCer-ThuE$^{283A}$-Aph$^{V224L}$ | FLIPsuc-$\Delta 2^{sCer\text{-}Aph}$ with W283A substitution and V224L in Aphrodite | this study |
| pRSET_sCer-ThuE$^{283\}}$-Aph$^{V224L}$ | FLIPsuc-$\Delta 2^{sCer\text{-}Aph}$ with W283G substitution in ThuE and V224L in Aphrodite | this study |
| pRSET_sCer-ThuE$^{283C}$-Aph$^{V224L}$ | FLIPsuc-$\Delta 2^{sCer\text{-}Aph}$ with W283C substitution in ThuE and V224L in Aphrodite | this study |
| pRSET_sCer-ThuE$^{283I}$-Aph$^{V224L}$ | FLIPsuc-$\Delta 2^{sCer\text{-}Aph}$ with W283I substitution in ThuE and V224L in Aphrodite | this study |
| pRSET_sCer-ThuE$^{D192N}$-Aph | FLIPsuc-$\Delta 2^{sCer\text{-}Aph}$ with D192N substitution in ThuE | this study |
| pRSET_sCer-ThuE$^{D192N}$-Aph$^{S208F}$ | FLIPsuc-$\Delta 2^{sCer\text{-}AphS208F}$ with D192N substitution in ThuE and S208F in Aphrodite | this study |
| pRSET_sCer-ThuE$^{D192N}$-Aph$^{V224L}$ | FLIPsuc-$\Delta 2^{sCer\text{-}AphV224L}$ with D192N substitution in ThuE and V224L in Aphrodite | this study |
| pRSET_sCer-ThuE$^{D192N}$-Aph$^{S208F/V224L}$ | FLIPsuc-$\Delta 2^{sCer\text{-}AphS208F/V224L}$ with D192N substitution in ThuE and V224L and S208F in Aphrodite | this study |
| pRSET_sCer-ThuE$^{283A/D192N}$-Aph$^{V224L}$ | FLIPsuc-65$\mu\Delta 2^{sCer\text{-}sAph\ D192N}$ with D192N and W283A substitutions in ThuE and V224L in Aphrodite | this study |
| pRSET_sCer-ThuE$^{283G/D192N}$-Aph$^{V224L}$ | FLIPsuc-90$\mu$ $\Delta 2^{sCer\text{-}sAphD192N}$ with D192N and W283G substitutions in ThuE and V224L in Aphrodite | this study |
| pRSET_sCer-ThuE$^{283C/D192N}$-Aph$^{V224L}$ | FLIPsuc-450$\mu$ $\Delta 2^{sCer\text{-}sAphD192N}$ with D192N and W283C substitutions in ThuE and V224L in Aphrodite | this study |
| pRSET_sCer-ThuE$^{283I/D192N}$-Aph$^{V224L}$ | FLIPsuc-1.1m $\Delta 2^{sCer\text{-}sAphD192N}$ with D192N and W283I substitutions in ThuE and V224L in Aphrodite | this study |
| pRSET_sCer-ThuE$^{283V/D192N}$-Aph$^{V224L}$ | FLIPsuc-2.4m$\Delta 2^{sCer\text{-}sAphD192N}$ with D192N and W283V substitutions in ThuE and V224L in Aphrodite | this study |

**Table S4:** FLIPsuc-Δ2sCer-Aph affinity mutant series.

| **Name** | **Mutation in ThuE** | **Affinity ($\mu$M)** |
|---|---|---|
| FLIPsuc-100$\mu$Δ2$^{sCer-Aph}$ | W283A | 101 ± 0.01 |
| FLIPsuc-120$\mu$Δ2$^{sCer-Aph}$ | W283G | 121 ± 0.01 |
| FLIPsuc-550$\mu$Δ2$^{sCer-Aph}$ | W283C | 556 ± 0.03 |
| FLIPsuc-1.1mΔ2$^{sCer-Aph}$ | W283I | 1080 ± 90 |
| FLIPsuc-4.1mΔ2$^{sCer-Aph}$ | W283V | 4104 ± 0.37 |

**A**

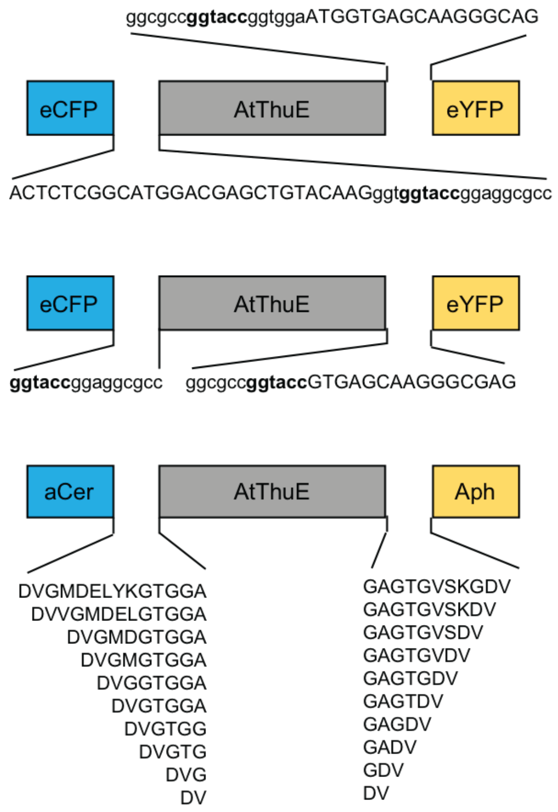


**B**

| | | | C-terminal linker | | | | | | | | | |
|---|---|---|---|---|---|---|---|---|---|---|---|---|
| | | | **R1** | **R2** | **R3** | **R4** | **R5** | **R6** | **R7** | **R8** | **R9** | **R10** |
| | | | DV | GDV | GADV | GAGDV | GAGTDV | GAGTGDV | GAGTGVDV | GAGTGVSDV | GAGTGVSKDV | GAGTGVSKGDV |
| N-terminal linker | **L1** | DV | -0.34 | -0.11 | -0.02 | -0.39 | -0.16 | -0.16 | -0.16 | -0.18 | -0.40 | -0.18 |
| | **L2** | DVG | -0.20 | -0.02 | -0.39 | -0.45 | -0.24 | -0.16 | -0.16 | -0.16 | -0.32 | -0.18 |
| | **L3** | DVGT | -0.20 | -0.02 | -0.47 | -0.59 | -0.31 | -0.19 | -0.04 | -0.23 | ND | -0.24 |
| | **L4** | DVGTG | -0.11 | -0.02 | -0.39 | -0.55 | -0.24 | -0.41 | -0.08 | -0.23 | -0.33 | 0 |
| | **L5** | DVGTGG | -0.08 | -0.06 | -0.54 | -0.61 | -0.13 | -0.61 | -0.14 | -0.26 | -0.37 | -0.32 |
| | **L6** | DVGTGGA | -0.06 | -0.06 | -0.53 | -0.58 | -0.29 | -0.61 | -0.08 | -0.23 | -0.26 | -0.43 |
| | **L7** | DVGGTGGA | -0.01 | -0.03 | -0.49 | -0.52 | -0.18 | -0.29 | -0.14 | -0.23 | -0.32 | -0.28 |
| | **L8** | DVGMGTGGA | -0.05 | -0.08 | -0.54 | -0.56 | -0.16 | -0.38 | -0.07 | -0.25 | -0.43 | -0.28 |
| | **L9** | DVGMDGTGGA | -0.05 | -0.06 | -0.65 | -0.57 | -0.28 | -0.67 | -0.49 | -0.32 | -0.42 | -0.41 |
| | **L10** | DVGMDEGTGGA | -0.05 | -0.21 | -0.69 | **-0.73** | -0.37 | -0.67 | -0.14 | -0.23 | -0.47 | -0.48 |
| | **L11** | DVGMDELGTGGA | -0.06 | -0.26 | -0.67 | -0.59 | -0.60 | -0.71 | -0.24 | -0.34 | -0.52 | -0.52 |
| | **L12** | DVGMDELYGTGGA | -0.07 | -0.26 | -0.50 | -0.73 | -0.43 | 0. | -0.61 | -0.20 | -0.52 | -0.58 |
| | **L13** | DVGMDELYKGTGGA | -0.01 | -0.22 | -0.57 | -0.46 | -0.42 | -0.24 | -0.61 | -0.46 | -0.49 | -0.50 |

**Figure S1. Emission ratio change is affected by modified linker length in FLIPsuc-Δ2$^{sCer-Aph}$**. Single amino acid truncation were performed on the linker residues connecting sCer and Aph to ThuE (A) , which altered *in vitro* emission ratio change of the nanosensors (B).
